# Prophages express a type IV pilus component to provide anti-phage defence

**DOI:** 10.1101/2024.03.29.587342

**Authors:** Kristina M. Sztanko, P. Nathael Javorčík, Alexa D. Fitzpatrick, Tatiana Lenskaia, Karen L. Maxwell, Alan R. Davidson

## Abstract

Phage genomes integrated within bacterial genomes, known as prophages, frequently encode proteins that provide defence against further phage infection. These proteins often function at the cell surface and prevent phages from attaching to their host receptor. Here, we describe the discovery of prophage-encoded proteins that resemble FimU, a component of the *Pseudomonas aeruginosa* type IV pilus. These phage FimU proteins are incorporated into the pilus without altering its function, yet they mediate robust protection against infection by phages that bind to the tip of the pilus, where FimU is likely located. The phage FimU proteins and the phage tail proteins that interact with FimU are highly diverse, suggesting that evolution in this system is driven by phage versus phage competition. To our knowledge, this is the first example of anti-phage defence mediated by replacement of a bacterial cell surface component with a phage-encoded protein.

## Introduction

Most bacterial genomes contain integrated phage genomes, which are known as prophages. While prophages can be induced and mediate formation of viral particles, their frequent persistence within bacterial genomes has led to symbiotic relationships between prophages and their bacterial hosts. An important manifestation of this symbiosis is that prophages generally express genes that provide a fitness advantage to their host^1^. These genes can enhance fitness by many means, frequently by providing defence against infection by other phages. The characterization of prophage-encoded anti-phage defences has led to the discovery of many novel and intriguing mechanisms to inhibit phage replication^2–8^.

One common way that prophages defend against phage infection is by preventing phage adsorption to the cell surface. For example, phages infecting *Pseudomonas aeruginosa* (*Pae*) almost invariably adsorb to cells by binding lipopolysaccharide (LPS) or the type IV pilus (T4P). The prophage of *Pae* phage D3 produces proteins that alter the LPS, which prevents adsorption by many LPS-dependent phages^9^. Similarly, several different *Pae* prophage-produced proteins have been identified that prevent normal T4P function so that phages requiring this structure for adsorption are unable to infect^2–4^.

Here, we focus on phages that bind to the T4P. The T4P is a long, thin structure that extends from the surface of the cell and is involved in a variety of processes including twitching motility^10^. The T4P is largely made up of the major pilin, PilA, which dynamically polymerizes and depolymerizes to extend and retract^10,11^. The T4P also contains five minor pilins (PilE, PilX, PilV PilW and FimU), which are found in lower abundance and are predicted to be located at the tip of the T4P^12–15^. Finally, Pae encodes a T4P adhesin protein, PilY1, which is thought to be involved in surface sensing and epithelial cell adhesion^16–18^. The minor pilin FimU, which is a focus of this study, is a component of the assembled T4P^12,14^, and interacts with the major pilin, PilA and the minor pilins PilE, PilV and PilX^15,19^. The minor pilin proteins have been proposed to form an initiation complex required for PilA polymerization. Specifically, FimU and PilE likely act as adaptors to connect the major pilin PilA to the remaining minor pilins and PilY1. In this model, the minor pilins are positioned at the tip of the T4P, with PilA monomers polymerizing underneath the minor pilin complex^19^. Although no structures of the complete T4P have yet been solved to prove this model, many different lines of evidence strongly support it^12,20–23^. The T4P mediates motility by adhering to surfaces through interactions with its tip, and then retracting the pilus structure back into the cell. Phages that bind the T4P are likely pulled to the cell surface upon T4P retraction^14^, but the subsequent steps leading to genome injection are not known.

This study investigates a group of T4P-dependent phages related to *Pae* phage JBD68, which we isolated in a previous study^5,25^. These phages, which we refer to as F10-like after the first described phage of this type^26,27^, are united by a very similar set of proteins that comprise their virion structures. They are distinguished from other characterized phages by possessing a unique set of proteins at their tail tip^17^. We were surprised to discover that the genomes of several F10-like phages encode a protein with significant sequence similarity to minor pilin protein FimU (∼30% pairwise identity across a 180 amino acid alignment). We hypothesized that this phage-encoded FimU homologue might serve to alter the function of the T4P as a means to protect F10-like prophage-containing cells from attack by other T4P-dependent phages.

In this paper, we show that the FimU proteins encoded in the genomes of F10-like phages do indeed alter the T4P to defend against phage infection. This defence operates within the context of prophage expression of these proteins, but functions only against F10-like phages, not other T4P-dependent phages. We have elucidated the mechanism of this anti-phage defence, and have also identified the phage proteins that likely interact with FimU to mediate cell surface attachment. This work illustrates a new mechanism for anti-phage defence and suggests that diversity among F10-like phages may be driven by phage versus phage competition.

## Results

### Phage encoded FimU-like proteins block infection by F10-like phages

Through analysis of the genome of F10-like *Pae* phage JBD68, sequenced as part of a previous study in our laboratory^5^, we discovered that this phage encodes a protein that is similar in sequence to the T4P component, FimU (28% identical to FimU of *Pae* strain PAO1). The gene encoding this phage FimU protein, referred to as P-FimU, lies within the cluster of genes encoding components of the phage tail, between the tail tube and the tail assembly chaperone (Fig. 1a). BLAST searches initiated using the JBD68 P-FimU protein as a query revealed that many prophages in *Pae* genomes encode P-FimU proteins. Genes encoding these proteins generally lay in the same genomic position as seen in JBD68, but other positions for these genes were also observed (Fig. 1a). The BLAST hits to the P-FimU proteins were found only in F10-like *Pae* prophages, not in other types of *Pae* prophages or prophages in species other than *Pae*. A representative alignment of all the phage encoded P-FimU proteins found in *Pae* showed that there are five distinct groups of these proteins (Extended Data Fig. 1a). We selected one representative of each of these groups for further study except for group 4, from which we selected both the JBD68 and Les3 proteins as we have both these phages in our collection and are used in our experiments described below. The selected P-FimU proteins were diverse in sequence with many pairwise percent identities below 40% (Fig. 1b,c).

**Fig. 1.**
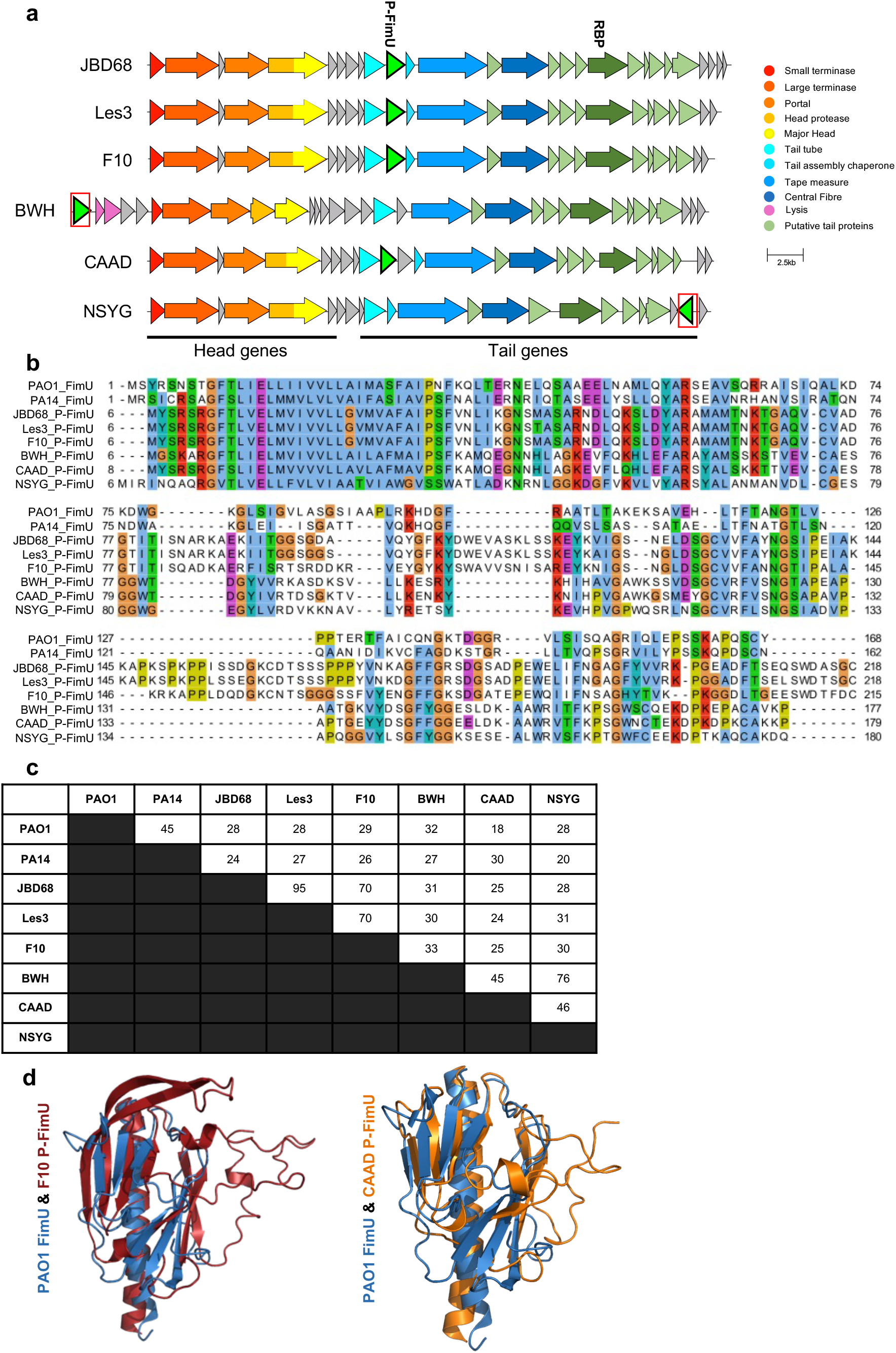
F10-like phages encode a protein similar to the FimU protein encoded by *Pae* strain PAO1. **a,** Genomic maps of the morphogenetic regions of F10-like phages are shown. Genes encoding universally required siphophage morphogenetic proteins are coloured as indicated in the legend. Putative tail proteins indicated in light green are homologous to proteins found in the JBD68 viral particle by mass spectrometry^25^. Genes encoding P-FimU proteins are indicated in bright green. Genes encoding P-FimU proteins found in atypical positions are boxed in red. **b,** Multiple sequence alignment of FimU and P-FimU proteins used in this study. **c,** Pairwise percent amino acid sequence identities among PAO1 and PA14 FimU and P-FimU proteins. **d,** On the left, a structural overlay of PAO1 FimU (blue) and the phage F10 P-FimU protein (red) is shown. On the right, PAO1 FimU (blue) and the CAAD P-FimU protein (orange) are overlaid. The P-FimU structures were predicted using AlphaFold2 and were truncated at the N-terminus to match the solved structure of PAO1 FimU (PBD: 4IPU).

To investigate whether the P-FimU proteins were similar in structure to *Pae* FimU protein, we used AlphaFold2 to generate predicted structures for each of the P-FimU proteins. We found that each of these structures overlaid well with the solved structure of *Pae* strain PAO1 FimU with root mean square deviations in structure ranging from 2.4 to 3.0 over at least 130 backbone positions (Extended Data Fig. 1b). The predicted P-FimU structures fell into 2 groups. The JBD68, F10, and Les3 structures formed one group, as expected from their similar sequences. The CAAD, BWH, and NYSG P-FimU proteins comprised the other group of similar structures despite displaying relatively low sequence similarity. Representative structures from each group are overlaid on the PAO1 FimU structure (Fig. 1d).

Since the F10-like phages and many others are dependent on the T4P for adsorption to the cell surface, we hypothesized that the P-FimU proteins might serve as an anti-phage defence by altering T4P formation. To address this issue, we assessed the effects of P-FimU protein expression on the replication of four different F10-like phages available in our collection: phages F10, JBD68, Les2 and Les3. Les2 and Les3 were identified as prophages of the hypervirulent strain LESB58^28,29^ and were induced and isolated from this strain. We also tested two MP22-like phages, DMS3 and D3112^5^. We expressed the six diverse P-FimU proteins described above (Fig. 1b,c) from plasmids in *Pae* strain PAK and assessed the effect of these proteins on the replication of these phages. Strain PAK was used for these assays because all of the phages were able to replicate on this strain. By spotting serial dilutions of lysates of each phage on strains expressing the P-FimU proteins, we found that each P-FimU protein completely blocked the replication of at least one of the four F10-like phages tested (Fig. 2a, b). Surprisingly, replication of phages DMS3 and D3112 was not affected by expression of any of the P-FimU proteins even though these phages require the T4P for replication. To further assess whether the effect of P-FimU proteins was specific to F10-like phages, we tested four other groups of previously studied, diverse T4P-dependent *Pae* phages. We found that all of them were unaffected by expression of P-FimU proteins (Extended Data Fig. 2). By contrast, expression of phage JBD26 Gp61, which was previously shown to block all T4P dependent phages^3^, prevented replication of all of the phages tested here (Fig. 2a, b & Extended Data Fig. 2). In summary, these assays showed that P-FimU proteins uniquely blocked F10-like phages, yet had no effect on the replication of five other distinct groups of T4P-dependent phages.

**Fig. 2.**
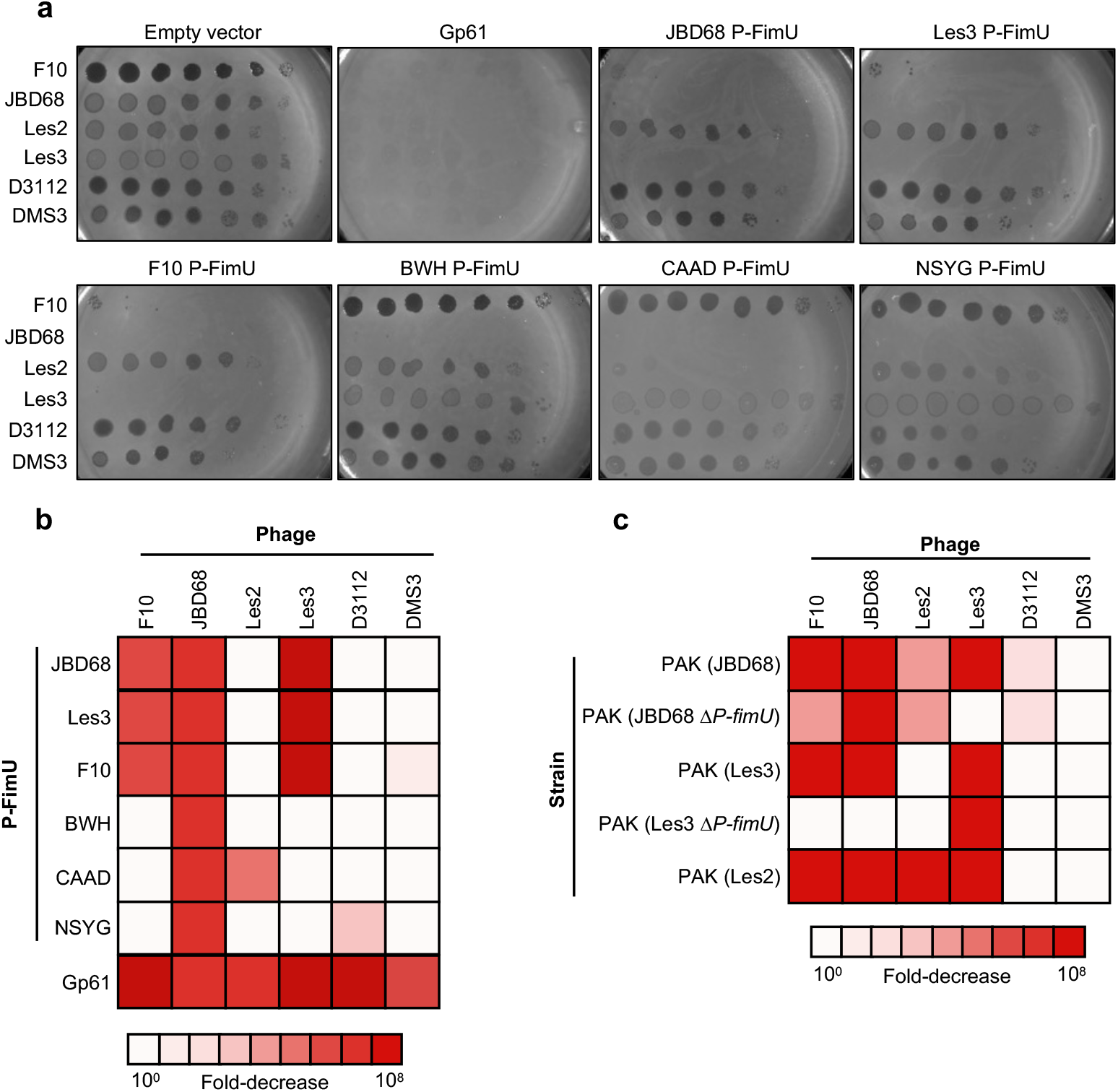
Expression of P-FimU proteins inhibits replication of F10-like phages. **a,** Ten-fold dilutions of lysates of F10-like phages and non-F10-like T4P-dependent phages (D3112 and DMS3) were spotted onto lawns of strain PAK expressing the indicated P-FimU protein or JBD26 Gp61 from a high copy number plasmid (pHERD30T). Arabinose was added to induce gene expression. Les2 P-FimU was not tested as it only differs in a single amino acid from Les3 (position 152). **b,** The fold-decrease in plaquing efficiency caused by expression of each P-FimU protein, as observed in Fig. 2a, is represented as a heat map. **c,** A heat map summarizing the fold-decrease in plaquing efficiency observed when the indicated phages were spotted onto PAK bearing JBD68 or Les3 prophages. Prophages harboring deletion mutations in the genes encoding P-FimU were also tested. The identity of the prophage tested is shown in parentheses.

Interestingly, expression of the different P-FimU proteins had variable effects on the F10-like phages tested. While the JBD68, F10 and Les3 P-FimU proteins, which are similar in sequence (Fig. 1b, c), all blocked phages F10, JBD68, and Les3, the BWH and NSYG P-FimU proteins blocked only phage JBD68. The CAAD P-FimU protein displayed a unique behaviour in being the only one that blocked phage Les2, yet it was unable to block phages F10 and Les3 (Fig. 2a, b). The expression of P-FimU proteins in *Pae* strain PAO1 resulted in an identical pattern of phage resistance (Extended Data Fig. 3). These results together suggest that F10-like prophages may express P-FimU proteins to defend against superinfecting phages.

### P-FimU proteins confer anti-phage defence when expressed from prophages

To determine whether expression of P-FimU proteins by prophages confers resistance to superinfecting phages, phage replication was tested on lysogenic strains bearing F10-like prophages. We found that these prophages did mediate resistance to other F10-like phages (Fig. 2c, Extended Data Fig. 4a). However, none blocked the replication of phages D3112 and DMS3, indicating that the T4P was still an available surface to act as a receptor for some phages. Phage Les2 was able to form plaques on JBD68 and Les3 lysogens, which is consistent with our observation that plasmid-expressed P-FimU proteins from these phages did not block Les2 replication (Fig. 2a, b). The partially attenuated replication of phage Les2 on the JBD68 lysogen is likely due to the expression of a phage JBD26 Gp61 homologue that is encoded in the JBD68 genome. We did not test a phage F10 lysogen in these experiments because this phage was not able to form lysogens in strain PAK.

To determine if the resistance mediated by the F10-like prophages was due to P-FimU protein expression, we generated deletions in the genes encoding these proteins in phages JBD68 and Les3. These deletions had no effect on the replication of these phages (Extended Data Fig. 4b). Lysogens bearing these mutant prophages were sensitive to all of the F10-like phages tested except when the infecting phage matched the prophage (Fig. 2c, Extended Data Fig. 4a). In these cases, replication was blocked by the prophage-expressed repressor protein, which is required to maintain prophages in an inactive state. The repressor proteins of the F10-like phages used here are diverse in sequence (Extended Data Fig. 5a), so repressor-mediated cross-immunity between these phages was not observed. These data imply that P-FimU proteins are expressed from prophages and that the level of expression is sufficient to block the replication of superinfecting phages.

Consistent with our conclusion that P-FimU proteins are expressed from prophage genomes, we found that the DNA regions upstream of five of the six genes encoding these proteins contained predicted promoters. Furthermore, the sequences of these upstream regions were highly conserved (Extended Data Fig. 5b) even though the downstream ORFs encode diverse P-FimU proteins. In the case of the gene encoding BWH P-FimU protein, conservation of the promoter region is still seen despite the different genomic context of this gene compared to the others. Given the high similarity of these promoter regions, we expect that all of these genes would be expressed from the prophages in a manner similar those of phages JBD68 and Les3. The upstream region of the gene encoding the NSYG P-FimU protein was not conserved; thus, we cannot predict its level of expression from the prophage.

### Phage encoded FimUs mediate normal T4P-dependent motility and phage infection

We hypothesized that the P-FimU proteins inhibit phage replication by incorporating into the T4P. To investigate this idea, we introduced the P-FimU expressing plasmids into a *Pae* PAO1 Δ*fimU* strain. Previous studies have shown that PAO1 Δ*fimU* strains are deficient in T4P-dependent activities, including twitching motility and phage adsorption. Using a standard twitching motility assay, we showed that the Δ*fimU* strains containing plasmids expressing the P-FimU proteins displayed similar levels of twitching motility as seen for the wild-type PAO1 strain, or the Δ*fimU* mutant bearing plasmids expressing bacterial FimU proteins from strain PAO1 or PA14 (Fig. 3a, Extended Data Fig. 6a). By contrast, the Δ*fimU* strain carrying empty vector was markedly reduced in its level of motility.

**Fig. 3.**
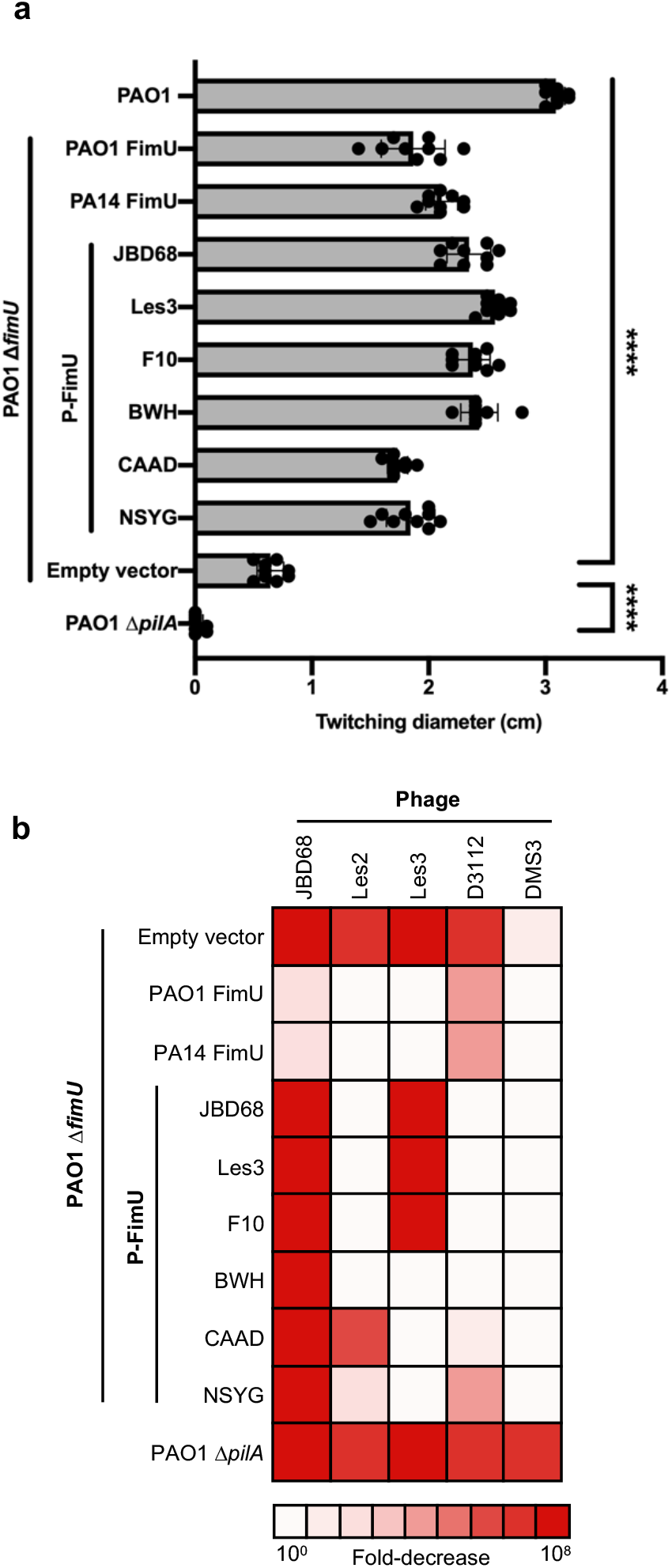
Expression of P-FimU proteins restores T4P activity in a PAO1 Δ*fimU* strain. **a,** The twitching diameters of PAO1 Δ*fimU* strains expressing the indicated P-FimU proteins. **** p < 0.0001 by one-way ANOVA using multiple comparisons relative to empty plasmid vector. **b,** Heat map representing the fold-decrease in plaquing efficiency relative to wild-type PAO1 of the indicated phages on lawns of a PAO1 Δ*fimU* strain expressing the indicated P-FimU proteins from a plasmid.

Expression of the P-FimU proteins fully restored the ability of the T4P-dependent phages, DMS3 and D3112, to replicate, further demonstrating that the P-FimU proteins are able to mediate normal functions of the T4P. The pattern of replication displayed by the F10-like phages on strains complemented by the P-FimU proteins mirrored the results seen when these proteins were expressed in the wild-type PAO1 or PAK strains (Fig. 3b, Extended Data Fig. 6b). As expected, plasmid-based expression of the *fimU* gene of strain PAO1 restored replication of all phages tested as did expression of the *fimU* gene from strain PA14, which encodes a protein that is 45% identical to the PAO1 homologue. It should be noted that phage DMS3 displays some ability to replicate in the PAO1Δ*fimU* strain though its replication increased by at least 10-fold in the complemented strains. This is may be due to the presence of shortened T4P in these strains, which also mediate a low level of twitching^14^. None of the phages tested were able to replicate on the Δ*pilA* strain, which lacks PilA, the major T4P subunit. The ability of Δ*fimU* strains complemented by the P-FimU proteins to both support T4P-dependent phage replication and wild-type levels of twitching motility strongly supports the conclusion that these proteins are incorporated into the T4P.

### JBD68 interacts with the tip of the T4P

Since the P-FimU proteins provide protection selectively against the F10-like phages, we surmised that these phages interact with the T4P in a fundamentally different manner. Previous studies using electron microscopy to investigate 12 different T4P-dependent *Pae* phages have shown that they bind along the length of the T4P, suggesting they interact with the major pilin protein, PilA (Supplementary Table 1). Protection against the F10-like phages provided by the expression of P-FimU proteins suggests that these phages might interact with bacterial FimU. To address this issue, we incubated phages JBD68 and DMS3 with strain PAO1 and visualized the phage-T4P interaction using negatively stained transmission electron microscopy. We found that phage DMS3, which is not blocked by P-FimU protein expression, bound to varying positions along the T4P fibre, which is comprised of the PilA protein (Fig. 4 a, c; Extended Data Fig. 7) By contrast, the F10-like phage, JBD68, exclusively bound to the tip of the T4P (Fig. 4 b, c; Extended Data Fig. 7). These results support our hypothesis that the F10-like phages bind to the T4P in a distinctive manner compared to other T4P-dependent phages. Furthermore, FimU and the other minor pilin protein components of the T4P are believed to be positioned at the tip^12–15,20–22^. Thus, these results suggest that F10-like phages interact with FimU at the tip of the T4P, and that the presence of P-FimU proteins at this position blocks phage adsorption. Notably, JBD68 is, to our knowledge, the only tailed phage shown to bind the tip of the T4P.

**Fig. 4.**
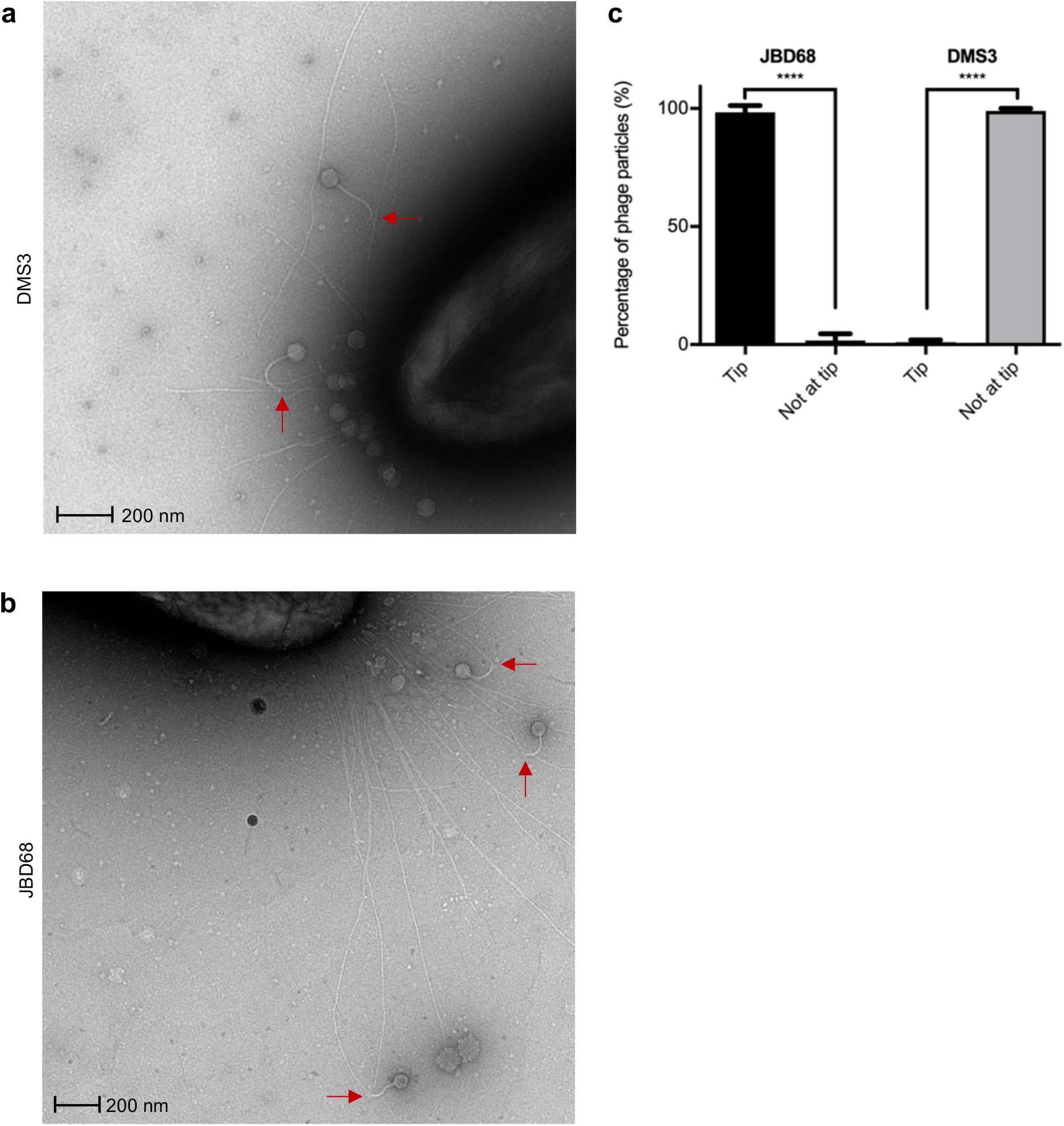
F10-like phages bind to the tip of the T4P. **a & b,** Negatively-stained transmission electron micrographs of phages DMS3 (**a**) and JBD68 (**b**), interacting with the strain PAO1 T4P. Interaction is indicated by red arrows. Images were taken at various magnifications to maximize the number of phages in the field of view. **c**, The percentage of JBD68 and DMS3 particles bound at different locations of the T4P is shown with mean and standard deviation across three independent replicate microscope grids. ****p < 0.0001 by one-way ANOVA using multiple comparisons between tip and not at tip.

### Identification of putative phage tail FimU-binding proteins

To gain further insight into the interaction of F10-like phages and the T4P, we sought to identify the tail tip proteins of these phages that may interact with FimU at the T4P tip. We reasoned that this protein in the F10-like phages tested here would vary in sequence as these phages display distinct patterns of inhibition in the presence of different P-FimU proteins (Fig. 2). The putative tail tip proteins of phage JBD68 were previously identified through genomic analysis and mass spectrometry of the JBD68 viral particle^25^. Comparison of the sequences of these putative tail tip proteins among the four F10-like phages used in this study showed that only two of these proteins varied markedly in sequence among these phages (Extended Data Fig. 8a). One of these proteins, the product of gene *24* in JBD68 (Gp24) was found to be non-essential when a deletion was created in the gene (Extended Data Fig. 8b). Thus, we focused on the other variable tail tip protein, which we refer to as the F10-like RBP (Receptor Binding Protein; Extended Data Fig. 8a, c).

To determine whether the F10-like RBPs were involved in the response to P-FimU-mediated inhibition of replication, we deleted the gene encoding this protein in a JBD68 prophage.

Induction of this mutant prophage resulted in the production of no infectious phage particles. We complemented this mutation by expressing the JBD68 RBP from a plasmid (referred to as pRBP_JBD68_) in the same strain (Fig. 5a). Phages produced by the complemented prophage replicated robustly on strain PAO1 containing a plasmid expressing pRBP_JBD68_. Performing the same test in a phage Les3 lysogen showed that the complemented phage was, like wild-type phage JBD68, unable to replicate due to the expression of the Les3 P-FimU protein from the prophage (Fig.5b). We then complemented the JBD68Δ*rbp* phage with a plasmid expressing the RBP homologue from phage Les2 (RBP_Les2_). Remarkably, this complemented phage replicated robustly on the Les3 lysogen containing a plasmid expressing RBP_Les2_. Since phage Les2 is not inhibited by P-FimU expressed from the Les3 prophage, these results imply that this resistance to inhibition is mediated by RBP_Les2_. Collectively, these data suggest that these RBPs are the tail proteins responsible for the interaction with the T4P FimU protein.

**Fig. 5.**
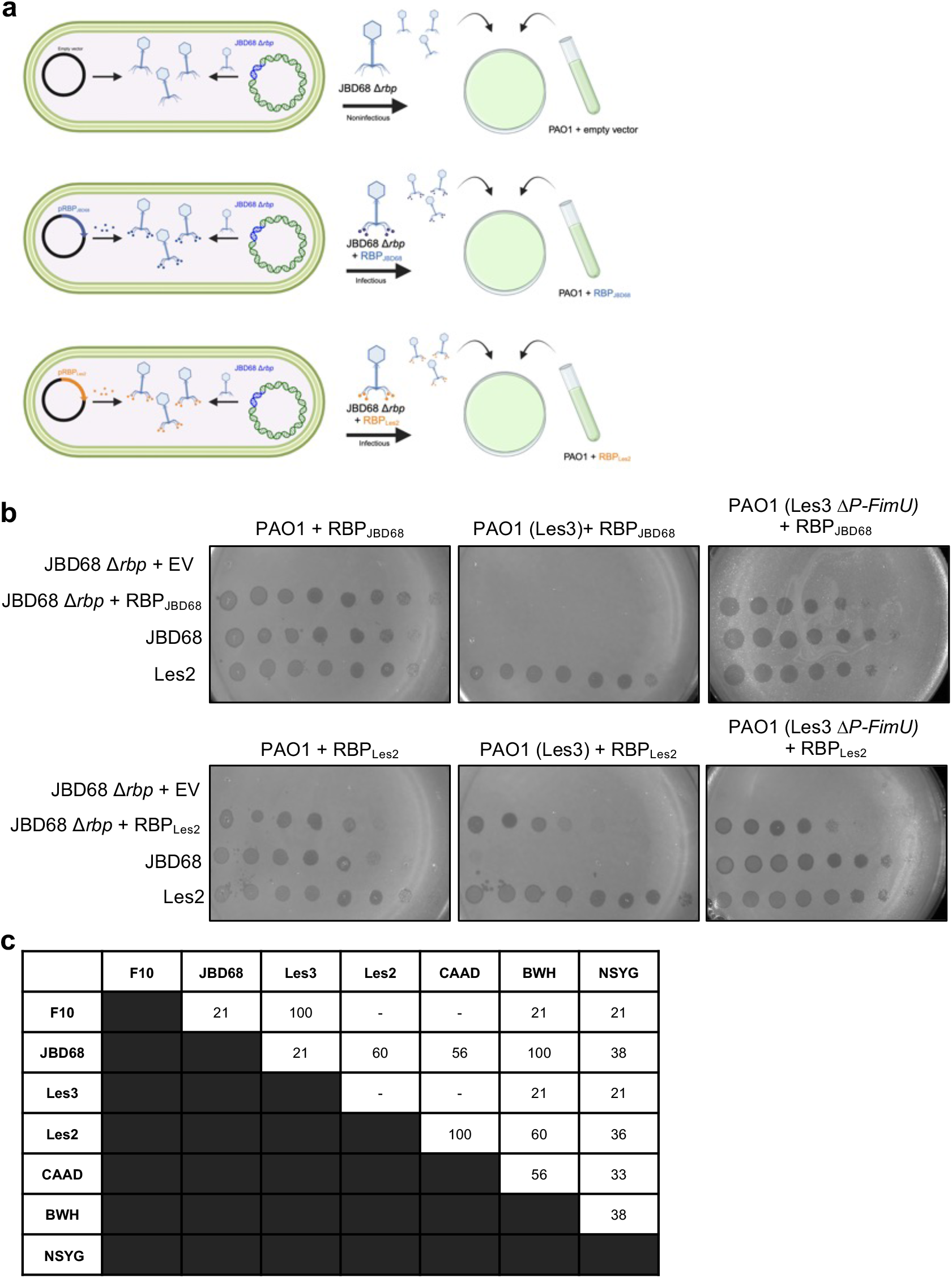
Swapping F10-like RBPs overcomes inhibition by P-FimU expressed from the Les3 prophage. **a,** A schematic of the JBD68 Δ*rbp* complementation experiment is shown. JBD68 Δ*rbp* prophages were induced in cells containing a plasmid containing empty vector, or plasmids expressing RBP_JBD68_ or RBP_Les2_. The resulting lysate was then plated on cells containing the plasmids expressing RBP_JBD68_ or RBP_Les2_. **b,** Ten-fold dilutions of lysates of phage JBD68 Δ*rbp* complemented with the indicated RBP proteins were spotted onto lawns of the indicated strains. The prophages contained in these strains are indicated in parentheses. The indicated RBP proteins were expressed from a plasmid. **c,** Pairwise percent amino acid sequence identities of RBPs from the indicated phages are shown. “–” denotes no identity.

The identity of the RBPs as the FimU-interacting proteins is supported by the pattern of phage replication inhibition by the P-FimU proteins (Fig. 2a, b) as compared to the pairwise sequence identity among the RBPs (Fig. 5c). Phages F10 and Les3 possess identical RBPs and display identical patterns of inhibition by the P-FimU proteins. Both phages are inhibited by the JBD68, Les3, and F10 P-FimU proteins, but not by those from phages BWH, CAAD, and NSYG. The patterns of inhibition of phages JBD68 and Les2 are each distinct from the other tested phages, and the sequences of their RBPs are also unique among these phages.

### Other phage genomes encode proteins resembling T4P structural components

Given the effective anti-phage defence provided by the P-FimU proteins, we wondered whether other phages might use similar mechanisms to block phage replication. To address this question, we searched for T4P structural components encoded in other phage and prophage genomes. We performed BLAST searches on a database of phage and bacterial genomes established in our laboratory in which genes encoding phage-related proteins have been thoroughly annotated^30^. Although this database contains only approximately 750 tailed phage genomes and 2200 bacterial genomes, we were able to identify one additional example of a T4P component encoded in prophages. In this case, proteins related to the major T4P subunit, PilA, were encoded in three distinct types of prophages (myophage, siphophage, and podophage) in five different strains of *Xylella fastidiosa* (Extended Data Fig. 9a, b). The sequences of these P-PilA proteins fell into two groups that are approximately 50% identical to each other and to the *X. fastidiosa* PilA protein. Predicted structures of these proteins generated by AlphaFold2 could be overlaid on the predicted structure of the *X. fastidiosa* PilA protein with RMSD values below 3 Å (Extended Data Fig. 9c).

We also searched the current NCBI database of phage genomes for proteins that resembled T4P components. We found four groups of proteins with sequence similarity to PilA encoded in the genomes of four different types of phages (Supplementary Table 2). These proteins may function similarly to the P-FimU proteins by incorporating into the T4P and inhibiting phage adsorption. One of these protein families appears to be truncated versions of PilA comprised of only the N-terminal 60 residues. This is very unlikely to be a sequencing error as similar proteins occur in six different *Pseudomonas* phages. During assembly, the conserved N-terminal alpha helix of PilA monomers are stacked and bundled together to form a hydrophobic core while variable C-terminal domains are surface exposed^23^. These shorter P-PilA proteins may still be able to incorporate into the T4P but would lack sites of phage binding. Alternatively, it is possible that they may inhibit T4P assembly and prevent T4P-dependent phage infection in this manner. Interestingly, none of the phages encoding the PilA-like proteins are temperate. However, it is well established that virulent phages, such as *E. coli* phage T4, express proteins early in infection that render the infected cells resistant to further superinfection^31–33^. These proteins are presumed to provide an advantage by preventing additional phages from adsorbing to cells that are already infected by the same phage.

## Discussion

We have shown here that F10-like prophages produce variant forms of FimU that can replace the normal function of *Pae* FimU while blocking the replication of related T4P-dependent phages. These proteins appear to act as molecular mimics, replacing the bacterial FimU proteins with P-FimU proteins that cannot be bound by phages. To our knowledge, this is the first example of phages replacing a bacterial cell surface component with a functional version that is phage resistant. As such, the P-FimU proteins represent a new type of prophage-encoded anti-phage system.

As described in the introduction, many lines of evidence imply that FimU and the other minor pilins are positioned at the tip of the pilus^12–15,20–22^. Since the replication of F10-like phages is blocked by FimU-like proteins, a parsimonious explanation for our observations is that the primary protein receptor for these phages is FimU and that the surface bound by these phages is absent or inaccessible in the P-FimU proteins. The use of FimU as a receptor by F10-like phages is well supported by our EM experiments showing that phage JBD68 binds exclusively to the pilus tip, a stark contrast to phage DMS3 that binds to the shaft of the T4P composed of PilA (Fig. 4). These EM results are consistent with our finding that the P-FimU proteins do not inhibit the replication of phage DMS3. In searching the literature, we did not find another example of a tailed phage that binds to the tip of a T4P as does JBD68, though 12 other *Pae tailed* phage have been shown by EM to bind the side of the T4P (Supplementary table 1). Our observation that JBD68 binds exclusively to the T4P tip further supports the conclusion that the T4P tip is composed of a distinct set of proteins that includes FimU.

Our demonstration that expression of the P-FimU proteins complemented *ΔfimU Pae* strains with respect to twitching motility implies that these phage-encoded proteins are incorporated into the T4P in a manner similar to the bacterial FimU protein (Fig.3a). Thus, our model for the mechanism of P-FimU function is that these proteins are incorporated into the T4P to impart normal function, but that they cannot be bound by F10-like phages. The AlphaFold2 predicted structures of the P-FimU proteins show that compared to the bacterial FimU proteins, they possess large loops and an extended C-terminus. These additional regions may occlude the surfaces required for phage binding (Fig. 1b, c). Interestingly, the F10, JBD68, and Les3 P-FimU proteins, which inhibit the most phages (Fig. 2a, b), possess two of these loop regions while the other three P-FimU proteins possess only one.

The P-FimU proteins are strikingly diverse with pairwise identities ranging down to 20%. These differences in P-FimU sequence are reflected by their varying inhibitory activities against the tested F10-like phages (Fig. 1, 2). In contrast to the P-FimU proteins, the sequences of FimU proteins encoded in *Pae* strains display little variation. BLAST searches revealed only two variants of FimU encoded in *Pae* strains, one matching FimU found in PAO1, and the other matching PA14 (45% sequence identity). Both of these *Pae* FimU proteins support the replication of all of the tested F10-like phages (Fig. 3b). Whether an F10-like phage will be inhibited by a P-FimU protein is determined by its RBP, which implies that these F10-like RBPs interact with FimU. These RBPs are also highly diverse in sequence at a level similar to the P-FimU proteins (Fig. 5c). The high sequence variability of the F10-like RBPs contrasts with the low variability seen in the *Pae* FimU proteins, which are likely the normal binding partners of these proteins. One explanation for this incongruity is that the diversity of the F10-like RBPs is a result of the challenge posed by the diverse P-FimU proteins. In this way, this aspect of the evolution of these phages may be driven much more by phage versus phage competition rather than phage versus host. In contrast with FimU, the sequences of PilA proteins across *Pae* strains are much more diverse^34^, likely a result of this major T4P subunit being targeted by many diverse phages. In this case, PilA diversity appears to be driven by direct phage-host competition. It should be noted that F10-like phages may bind to FimU and PilA as these two proteins interact at the tip of the T4P^15^. Overall, it appears that the F10-like phage precursors may have gained an initial evolutionary advantage by targeting FimU, which is not targeted by other phages. However, this new niche was subsequently narrowed by interphage competition mediated by the P-FimU proteins.

In summary, this work elucidates a mechanism by which phages express a phage receptor component as a means to block subsequent phage infections. Although this is the first described example of such a mechanism, our identification of T4P components encoded in other diverse phages suggests that this may be a widely employed method of anti-phage defense.

## Data availability

The data are available in the manuscript.

## Acknowledgments

The authors thanks members of the Davidson, Maxwell and Burrows (McMaster University, Hamilton, Ontario, Canada) laboratories for their helpful discussions. This study was supported by grants from the Canadian Institutes of Health Research to K.L.M. (PJT-165936) and A.R.D. (FDN-15427), and a Natural Sciences and Engineering Research Council Arthur B. McDonald Fellowship to K.L.M. (SMFSU-581368-2023 ). A.R.D. is the Canada Research Chair in Bacteriophage-Based Technologies.

## Author contributions

K.M.S. and A.R.D. conceptualized this work. K.M.S performed experiments with assistance from P.N.J and A.D.F.; K.M.S. and A.R.D. analyzed the data. Bioinformatics were performed by K.M.S., T.L. and A.R.D.. K.M.S and A.R.D. wrote this manuscript with input from K.L.M. All authors approve the final version and conclusions.

## Materials and Methods

### Bioinformatics

Phage encoded pilus proteins were identified using BLAST^35^ searches. Clinker^36^ was used to generate genomic maps of F10-like phages and all other prophages in this study. Multiple sequence alignments were generated in Jalview Version 2^37^ using MUSCLE^38^ with default settings. Pairwise alignment percent identities were also generated using Jalview Version 2. Predicted structures were generated using AlphaFold2^39^, proteins overlays were generated using Pymol 2.5.3^40^ and RMSD scores were generated using DALI^41^. Fig. 5a was created using BioRender.com (agreement number XT26KNL13O). Promoters were predicted using Promoter Calculator algorithm by De Nova DNA^42^. A list of all relevant accession numbers can be found in Supplementary Table 3.

### Bacterial strains and plasmids

Strains, phages and plasmids used in this work are listed in Supplementary Table 4. Strains were grown in 3 ml or 5 ml of Lysogeny broth (LB) or on 1.5% LB agar plates for 18 hr (overnight) at 37 °C unless otherwise stated. Plasmids were transformed into *P. aeruginosa* using electroporation and chemically competent *Escherichia coli* by heat shock as previously described^43^. When required, media was supplemented with 50 μg/ml or 30 μg/ml gentamycin for *P. aeruginosa* and *E. coli* respectively. Expression from the pBAD promoter on high copy number plasmid, pHERD30T, was induced with 5 mM L-arabinose. The type I-C CRISPR-Cas system expressed in strain PAO1 was induced with 0.5 mM isopropylβ-D-1 thiogalactopyranoside (IPTG).

### DNA cloning and manipulation

Lysogens of JBD68, F10 and Les3 were used for amplification of *p-fimU* genes using PCR. BWH, CAAD & NSYG *P-fimU* genes were synthesized by Twist Bioscience, South San Francisco, California, USA. All genes were cloned into pHERD30T using the restriction enzymes, *Nco*I & *Hind*III). Spacers targeting Les3 and JBD68 *p-fimU* were cloned into pHERD30T using the restriction enzyme BsaI as previously described^44^. All cloning and mutagenesis was confirmed via sequencing performed by The Centre for Applied Genomics, The Hospital for Sick Children, Toronto, Canada.

Small deletions in the phage genes encoding P-FimU proteins were created using a I-C CRISPR-Cas system bearing a helicase-attenuated mutant Cas3 protein, as previously described^44,45^. Strain PAO1 harbouring the mutant I-C CRISPR system was grown to OD_600_ = 0.4 at 37 °C. Phage was added at an MOI of 10 and grown at 37 °C for 18 hr. Phage lysates were collected as described below. *P-fimU* mutants were isolated by performing phage plaquing assays (described below) on PAO1 with an integrated I-C CRISPR system containing the wild-type helicase to facilitate efficient selection against wild-type phage. The resulting plaques were isolated in SM buffer and mutations were confirmed via PCR and DNA sequencing. Deletions in the JBD68 gene encoding Gp24 were made in the same manner.

A deletion mutant of JBD68 *rbp* was generated using Pex18Gm as previously described^46^. 500 nucleotide homology arms were used to facilitate an in-frame, internal deletion of 900 nucleotides (amino acids Δ80-380) of JBD68 *rbp* that would produce a non-functional protein, while maintaining potential operon expression downstream. Mutants were confirmed via PCR and DNA sequencing. All bacterial strains and oligonucleotides used in this study are summarized in Supplementary Tables 4 and 5.

### Phage lysate production and lysogen formation

Phage lysates were produced by growing lysogens at 37 °C overnight, allowing spontaneous prophage production. Overnight cultures were centrifuged for 2 min at 10,000 X *g* and the supernatant was collected. A few drops of chloroform were added to the supernatant and incubated for 30 min at room temperature, with rocking. Lysates were centrifuged one final time (10,000 X *g*, 10 min) to remove cell debris and stored at 4 °C. Phage lysates were normalized to 10^8^-10^10^ PFU/ml.

Lysogens were isolated by plating 10-fold serial dilutions of phage lysate on either PAO1 or PAK. Cells from the centre of zones of clearing streaked to single colonies and then streaked across their respective lysates to identify resistant colonies. Resistant colonies were then confirmed as lysogens by growing the putative lysogens at 37 °C overnight, collecting the supernatant, and plating the lysates on their respective host strains to confirm infectious phage particles were produced.

### Phage plaquing assays

Phage plaquing assays were performed as previously described with minor modifications^43^. 200 μl of overnight *P. aeruginosa* culture was added to 3 ml of 0.6% LB agar and top plated on a 1.5% agar plate containing 10 mM MgSO_4_ and, when necessary, 50 μg/ml gentamycin and 5 mM arabinose (expression from pHERD30T). Phage lysates were tenfold serial diluted in SM buffer (100 mM NaCl, 8 mM MgSO_4_ & 50 mM Tris-HCl (pH 7.5)) and spotted on the surface. Plates were incubated at 30 °C overnight. All spotting assays were repeated at least 3 times with high reproducibility. Data shown in figures depict one representative experiment.

### Twitching motility assays

Twitching motility assays were performed as previously described^47^ with the following modifications. Individual colonies were stabbed into 1% LB agar (1% agar, 1% tryptone powder, 0.5% yeast extract & 0.5% NaCl) supplemented with gentamycin and arabinose and poured in pre-treated tissue culture-grade plates (VWR). Three replicates were performed with a minimum of three colonies per replicate. Plates were incubated at 37 °C for 24 hours, inverted. The agar was then removed, and the plates were stained with 1% crystal violet for 5 min while shaking. Excess stain was washed away with distilled water. Stained twitching areas were measured manually and Ordinary One-way ANOVA with multiple comparisons was performed using GraphPad Prism version 9.0.1 for Mac, GraphPad Software, Boston, Massachusetts USA, www.graphpad.com.

### Transmission electron microscopy

Samples were prepared by resuspending a single colony of PAO1Δ*pilT* (this mutant strain was used because increased numbers of T4P accumulate on the surface of this strain ) in 20 μl of SM buffer. 10 μl of cell mixture was incubated with approximately 10^8^ phages for 30 minutes at room temperature. Alternatively, overnight cultures of PAO1 Δ*pilT* were subcultured and grown to approximately OD_600_ = 0.7. Cells were diluted ten-fold in SM buffer and 10 μl of this mixture were incubated with approximately 10^8^ phages for 30 minutes at room temperature. Samples were allowed to bind to glow-discharged carbon grids for 2 minutes. Samples were washed twice with double distilled water and stained with 2% uranyl acetate for 15 seconds. Dried grids were imaged on Talos L120C transmission electron microscope (Microscopy Imaging Laboratory, Faculty of Medicine, University of Toronto, Toronto, Canada). Phages bound at tip versus not at tip were counted (minimum 100 particles per sample) and Ordinary One-way ANOVA with multiple comparisons was performed using GraphPad Prism version 9.0.1 for Mac, GraphPad Software, Boston, Massachusetts USA, www.graphpad.com.

### F10-like RBP Complementation assays

A single colony of PAO1 (JBD68 Δ*rbp*) harbouring pHERD30T expressing either RBP_JBD68_, RBP_Les2_ or empty vector were grown in 5ml of LB supplemented with gentamycin and arabinose overnight. The following day, 2 ml of lysate were prepared as described above. Tenfold serial dilutions of prepared lysates were spotted as described above on lawns of PAO1 harbouring the same pHERD30T vector used to complement JBD68 Δ*rbp.* Plates were incubated at 30 °C overnight. Note that sufficient complementation required expression of the upstream gene (JBD68 ARM70480.1 & ARM70481.1[RBP_JBD68_] and Les2 WP_010791962.1 & WP_010791961[RBP_Les2_]) as expression of just *rbp* from either phage did not allow for robust replication, likely due to a stoichiometric requirement between the two gene products.

