## Extended Data Figures for "Prophages express a type IV pilus component to provide anti-phage defence"

**a**

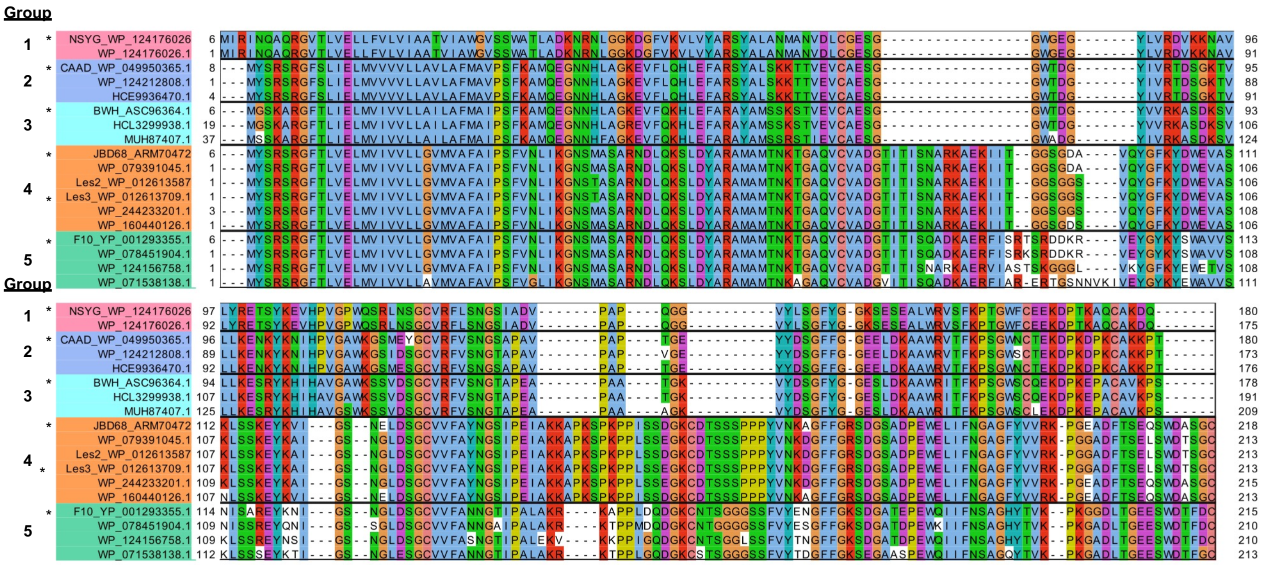

**b**

| | PAO1 | JBD68 | F10 | Les3 | BWH | CAAD | NSYG |
|---|---|---|---|---|---|---|---|
| **PAO1** | | 2.9 | 2.7 | 3 | 2.6 | 2.4 | 2.4 |
| **JBD68** | | | 2.3 | 2.6 | 2.7 | 2.6 | 2.4 |
| **F10** | | | | 2.2 | 2.4 | 2.4 | 2.1 |
| **Les3** | | | | | 3.3 | 3.4 | 2.7 |
| **BWH** | | | | | | 1.1 | 1.8 |
| **CAAD** | | | | | | | 1.4 |
| **NSYG** | | | | | | | |

**Extended Data Fig. 1| Identification and structural comparisons of P-FimU proteins in F10-like prophages. a,** A representative multiple sequence alignment of P-FimU proteins gathered from iterated PSI-BLAST searches using JBD68 P-FimU as a query. P-FimU protein groups are denoted by different colors. P-FimU proteins used in this study are indicated with asterisks. **b,** A comparison of the pairwise root mean square deviations in Ångstroms (RMSD) of the PAO1 FimU structure and the P-FimU predicted structures generated by Alphafold2.

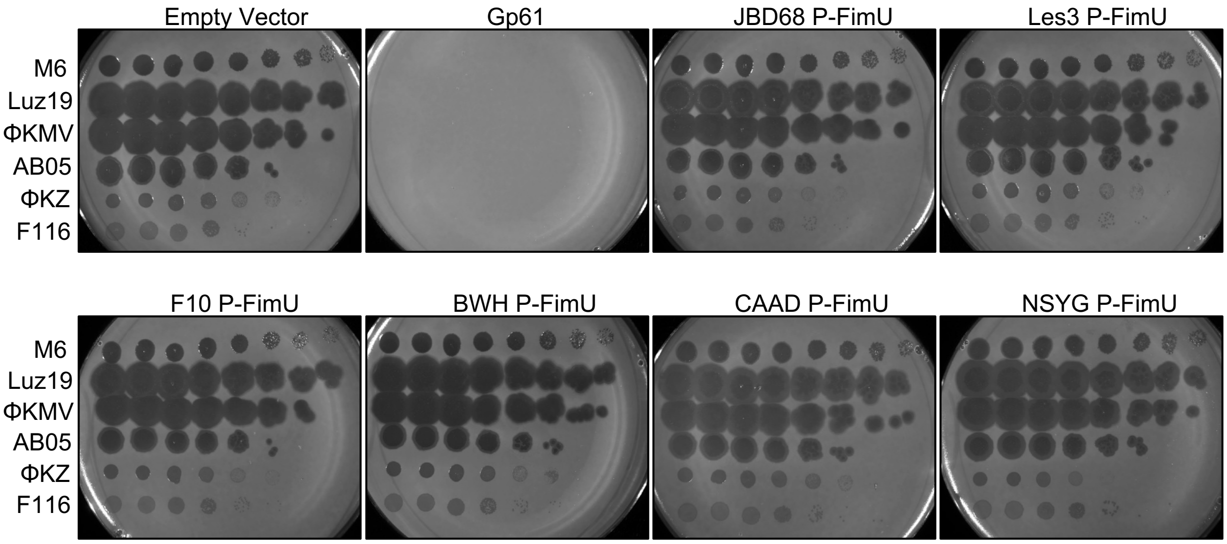


**Extended Data Fig. 2 | Expression of P-FimU proteins does not prevent infection of diverse T4P phages.** Ten-fold dilutions of the indicated phage lysates spotted on lawns of strain PAK expressing P-FimU proteins from plasmids. Distinct groups of T4P-dependent phages are composed of 1) M6, 2) Luz19, ΦKMV, AB05, 3) ΦKZ and 4) F116.

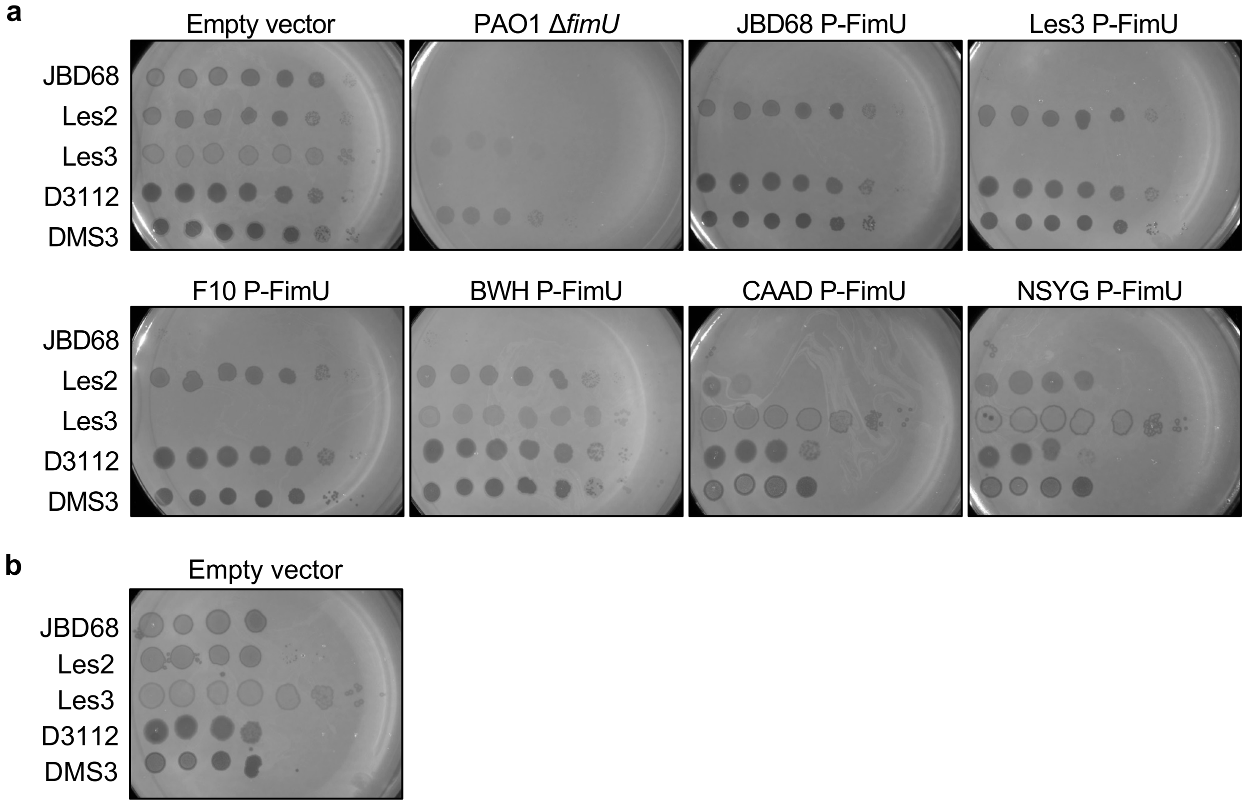


**Extended Data Fig. 3 | Expression of P-FimU proteins in strain PAO1. a,** Ten-fold dilutions of the indicated phage lysates spotted on lawns of strain PAO1 expressing P-FimU proteins from plasmids. Note that phage F10 was not tested on strain PAO1 as it does not replicate on this strain. Different lysate preparations were used for testing strain PAO1 expressing the CAAD and NSYG P-FimU. The results of spotting these lysates on strain PAO1 bearing an empty vector are shown in **b**.

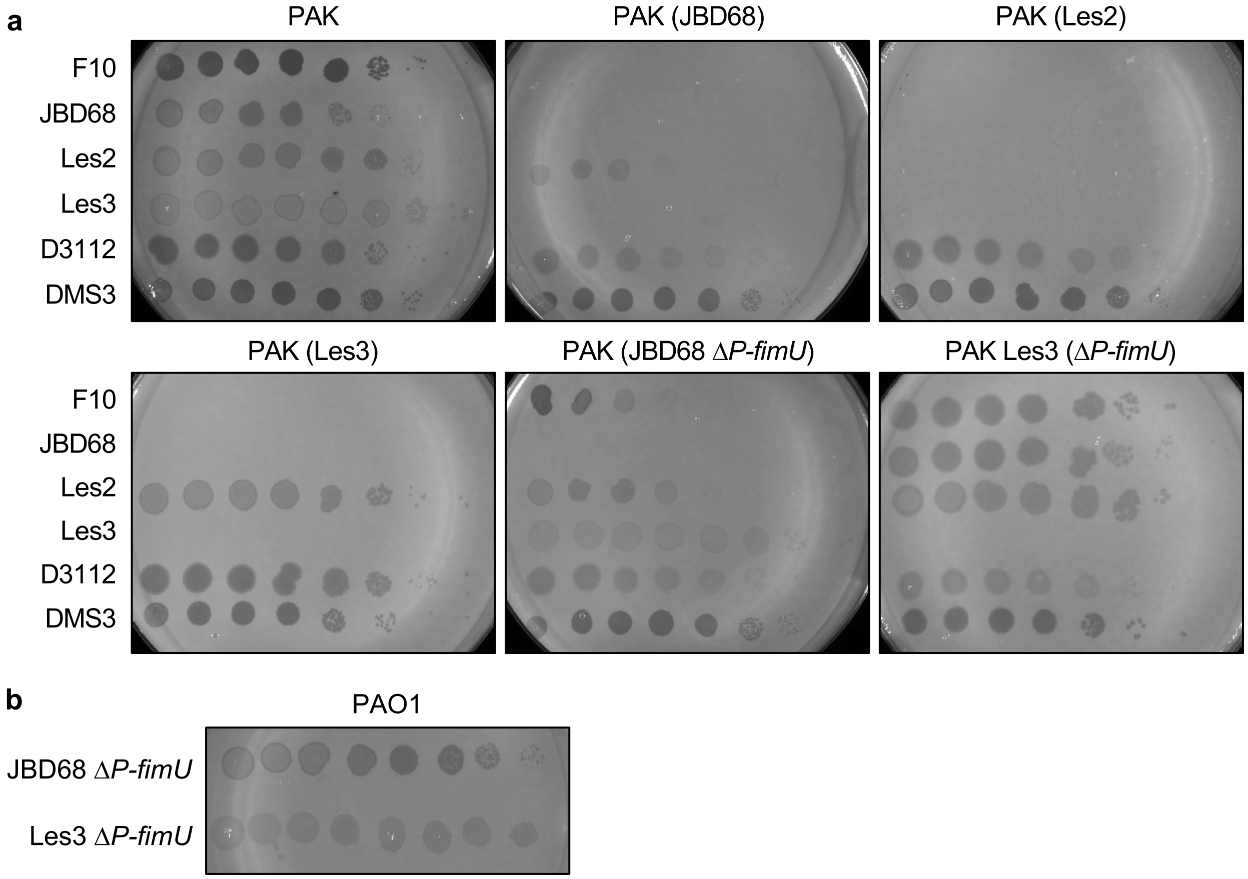


**Extended Data Fig. 4 | Mutations in the genes encoding P-FimU proteins in JBD68 and Les3. a,** Ten-fold dilutions of the indicated phage lysates spotted on lawns of strain PAK and PAK lysogens of wild-type JBD68 and Les3 as well as prophages harboring mutations in the genes encoding the P-FimU proteins. These results are displayed as a heat map in Fig. 2c. **b,** Ten-fold dilutions of lysates of JBD68Δ*p-fimU* and Les3Δ*p-fimU* mutants spotted on a lawn of strain PAO1.

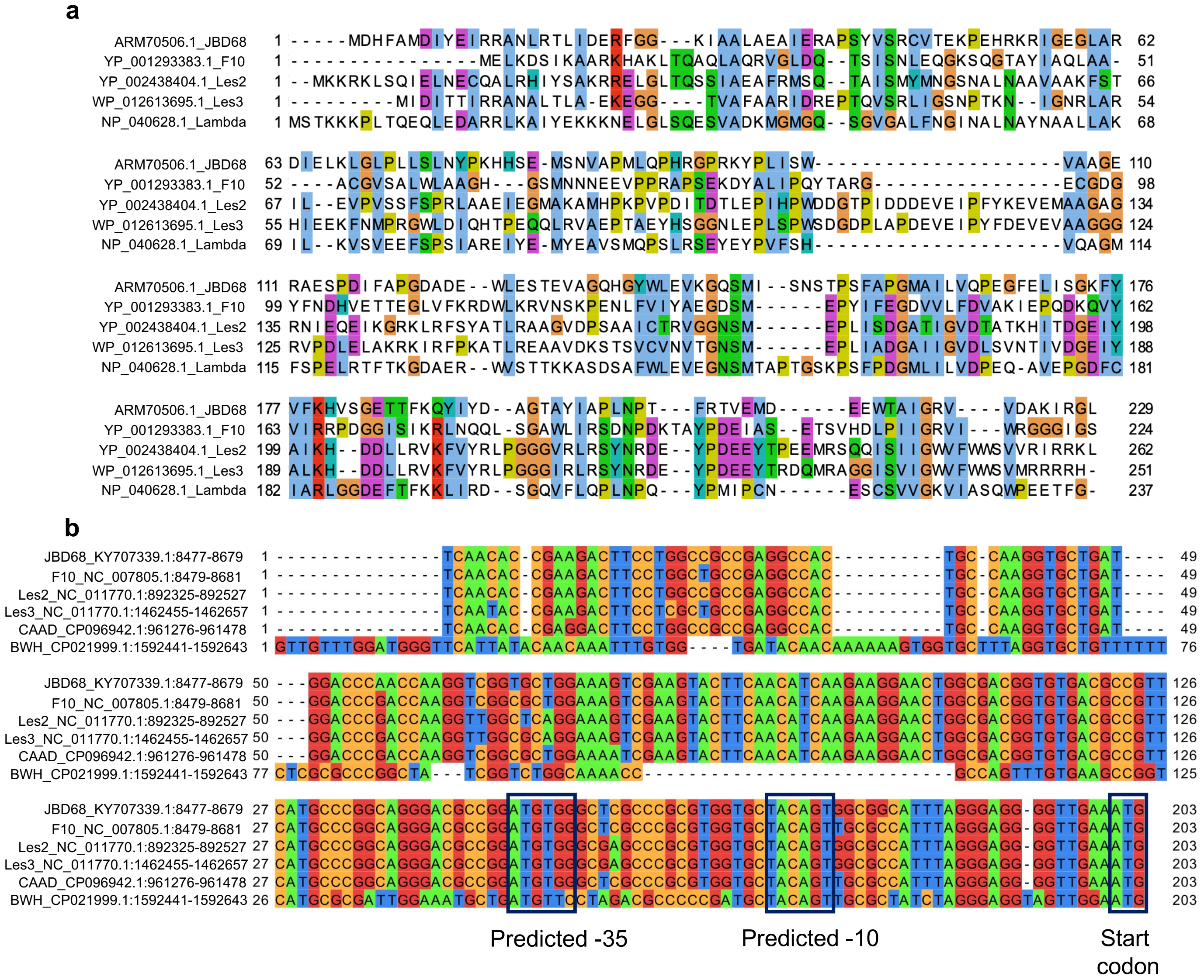


**Extended Data Fig. 5 | F10-like phage sequence alignments. a,** A multiple sequence alignment of repressor proteins of the F10-like phages. The well characterized *E. coli* phage lambda repressor is shown for comparison[1]. A higher degree of conservation is seen in the C-terminal domain, which mediates dimerization and self-cleavage. **b,** The putative promoter regions of P-FimU encoding genes. 200 nucleotides upstream of each P-FimU start codon are included. The -35 and -10 promoter sequences were predicted using the Promoter Calculator algorithm by De Nova DNA. A promoter was not detected for the NSYG P-FimU encoding gene and its upstream nucleotide sequence is highly diverged from the others.

a

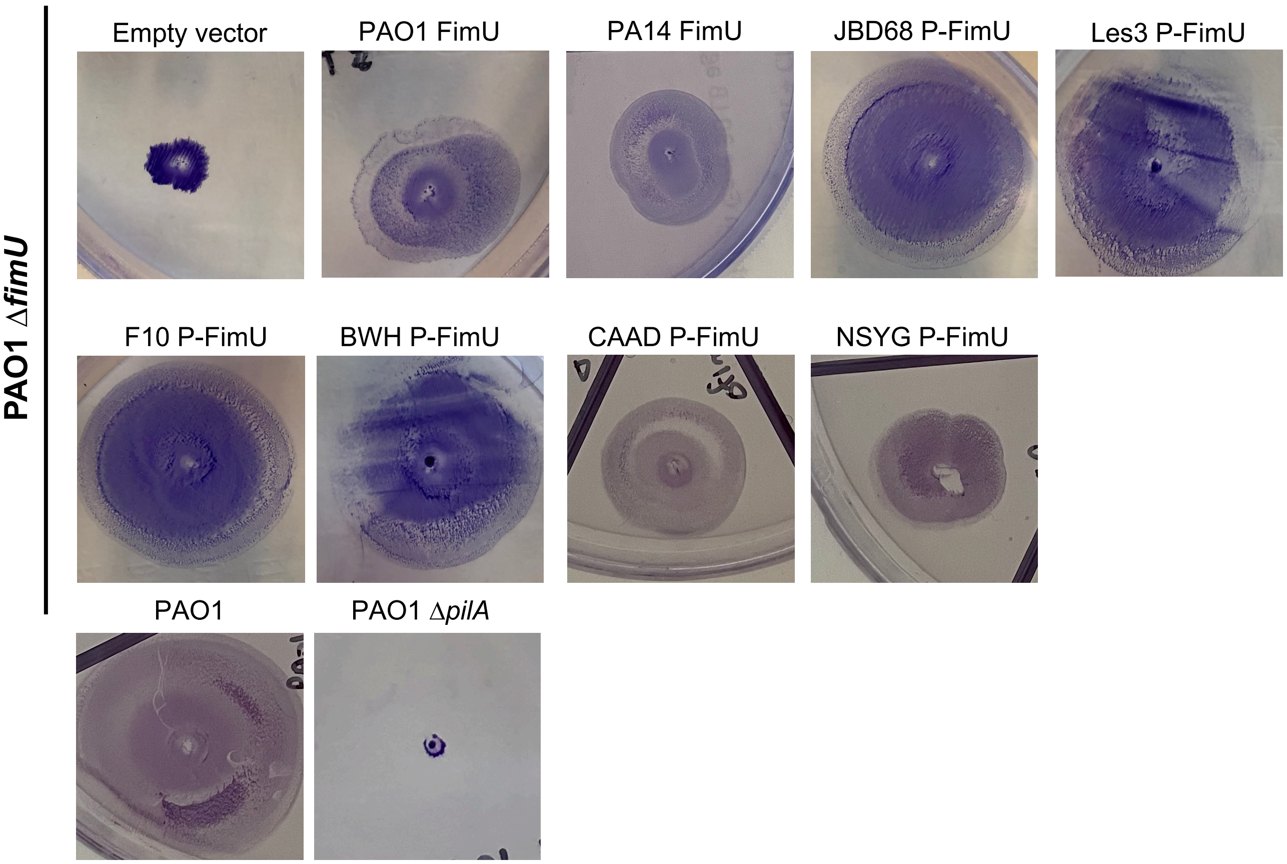

Empty vector
PAO1 FimU
PA14 FimU
JBD68 P-FimU
Les3 P-FimU
F10 P-FimU
BWH P-FimU
CAAD P-FimU
NSYG P-FimU
PAO1
PAO1 ΔpilA
PAO1 ΔfimU


b

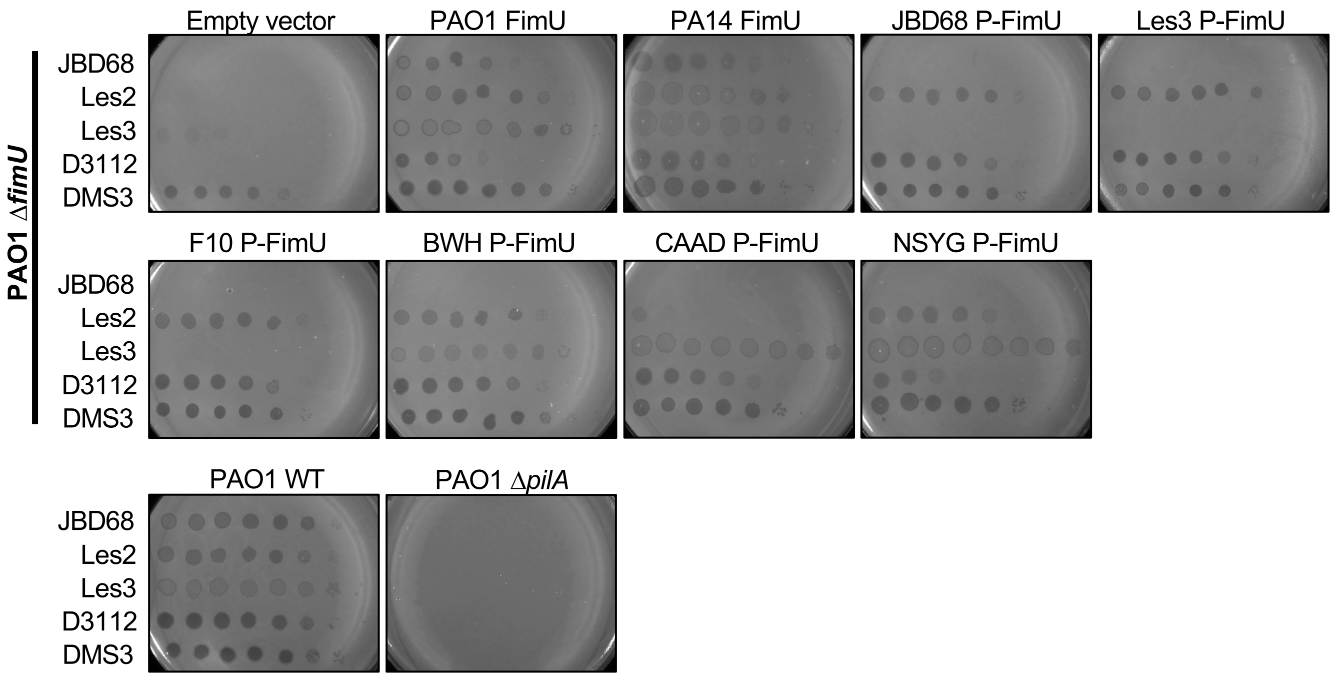

Empty vector
PAO1 FimU
PA14 FimU
JBD68 P-FimU
Les3 P-FimU
F10 P-FimU
BWH P-FimU
CAAD P-FimU
NSYG P-FimU
PAO1 WT
PAO1 ΔpilA
JBD68
Les2
Les3
D3112
DMS3
PAO1 ΔfimU

**Extended Data Fig. 6 | Expression of P-FimU proteins restores twitching motility in a PAO1Δ*fimU* strain. a,** Representative images of twitching motility assays of strain PAO1Δ*fimU* expressing different P-FimU proteins from a plasmid, as well as strains PAO1 wild-type and PAO1 Δ*pilA*. Twitching radii were stained with 1% wt/vol crystal violet. Averages of these experiments are shown in Fig. 3a. **b,** Ten-fold dilutions of lysates of the indicated phages spotted on lawns of strain PAO1 Δ*fimU* expressing P-FimU proteins from a plasmid. These results are displayed as a heat map in Fig. 3b

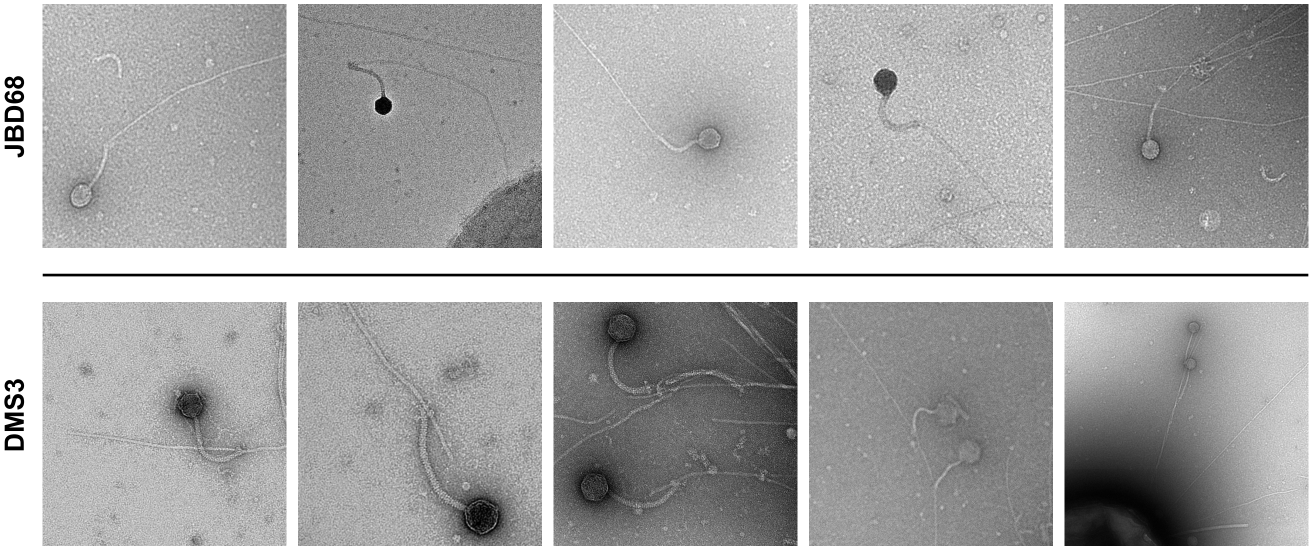


**Extended Data Fig. 7 | F10-like phages bind at the tip of the T4P**. Additional electron micrographs of phages JBD68 and DMS3, respectively, interacting with the PAO1 T4P. Images were taken at various magnifications to provide optimal fields of view.

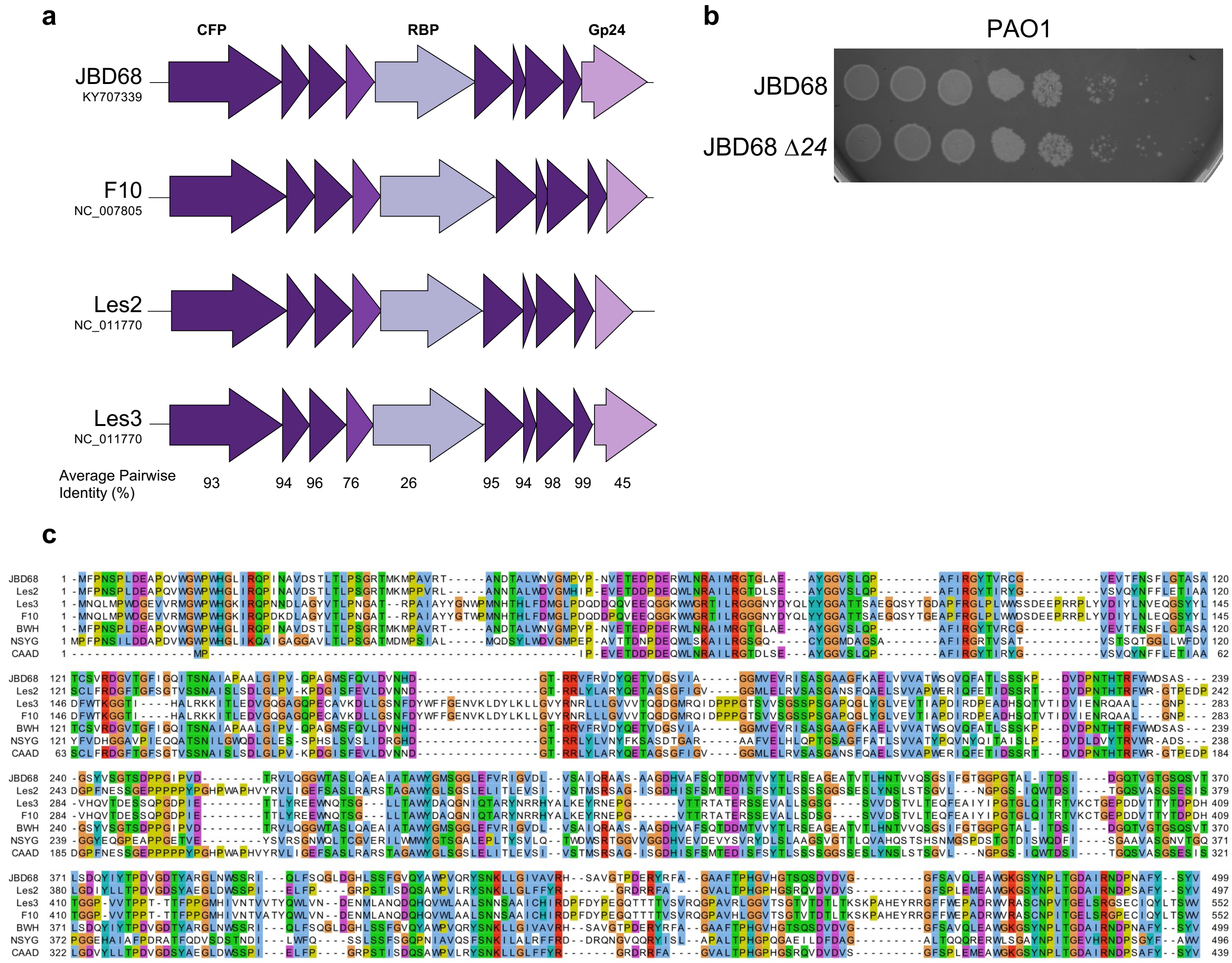


**Extended Data Fig. 8 | Identification of putative RBP in F10-like phages. a,** A schematic of the genes encoding the putative tail tip proteins of the four F10-like phages in our collection (CFP = central fibre protein). The average pairwise percent identity across all four phages is shown for each protein. **b,** Ten-fold dilutions of lysates of phage JBD68Δ*24* spotted on a lawn of strains PAO1. **c,** An amino acid multiple sequence alignment of RBP from F10-like phages is shown.

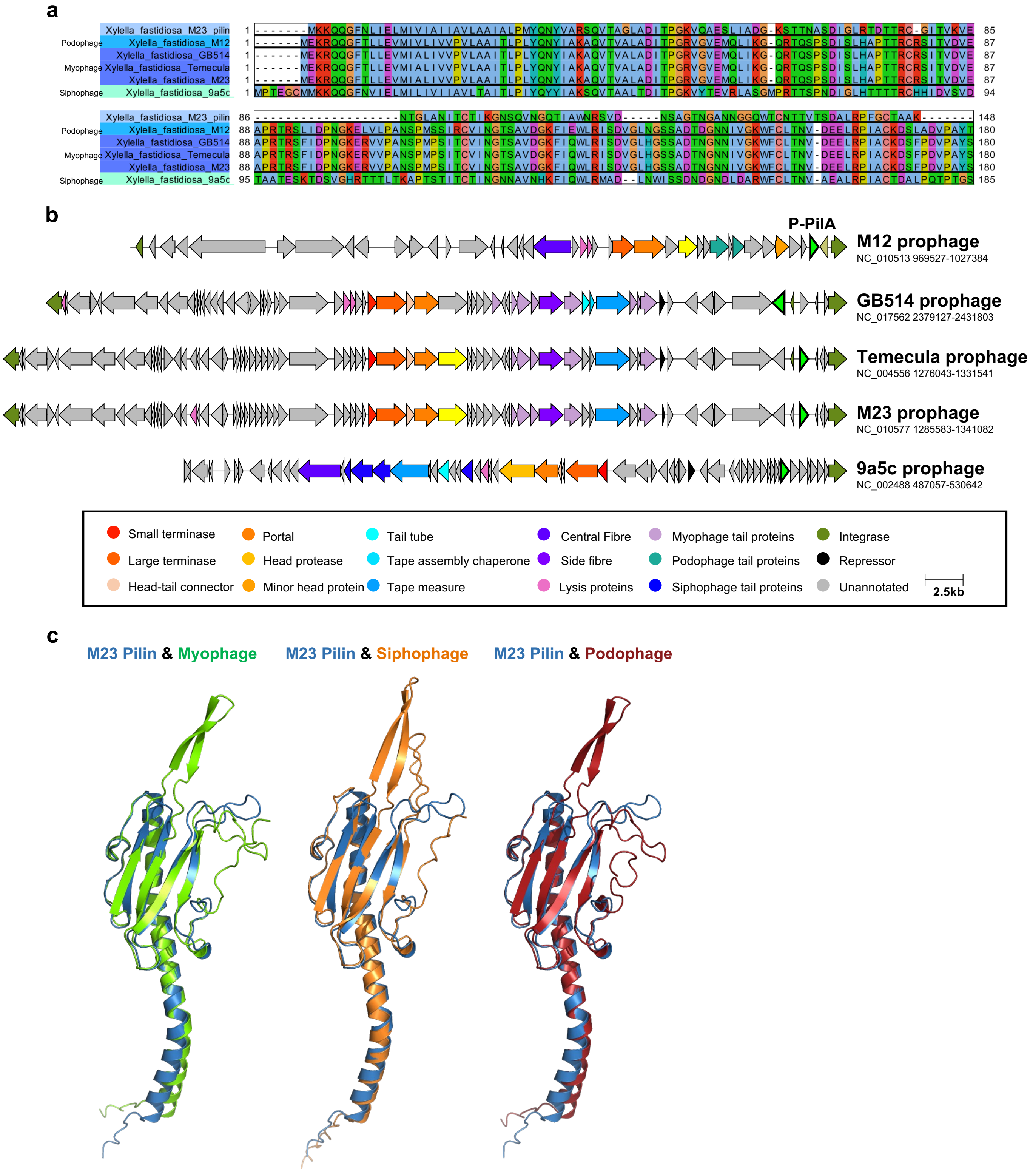


| | M23 Pilin | Myophage | Siphophage | Podophage |
|---|---|---|---|---|
| M23 Pilin | | 2.8 | 3 | 2.7 |
| Myophage | | | 2.1 | 0.9 |
| Siphophage | | | | 2.1 |
| Podophage | | | | |

**Extended Data Fig. 9 | Identification of T4P-like proteins in *Xylella fastidiosa* prophages. a,** Multiple sequence alignment of *X. fastidiosa* strain M23 pilin (PilA) and similar proteins encoded in five prophages. **b,** A schematic of the genomes of five *X. fastidiosa* prophages that encode a protein similar to the pilin protein of strain M23. Alignments were generated with Clinker and annotated function is denoted by color. Genes encoding pilin-like proteins are denoted in bright green **c,** Structural predictions and overlays of the M23 pilin (blue) and the pilin like proteins encoded by *X. fastidiosa* myo- (green), sipho- (orange) and podo- (red) prophages. Structures were predicted using AlphaFold. A comparison of the RMSD scores in Ångstroms of the M23 pilin and P-PilA predicted structures generated by Alphafold2.

**Extended date references**

1. Johnson, A., Poteete, A., Lauer, G. *et al.* λ Repressor and cro—components of an efficient molecular switch. *Nature* **294**, 217–223 (1981). https://doi.org/10.1038/294217a0
