## Supplementary Tables for "Prophages express a type IV pilus component to provide anti-phage defence"

| Phage | Location | Method | Host organism | Source |
|---|---|---|---|---|
| Ab05 (podophage) | T4P | T4P mutants | *Pae* | [1] |
| **C22 (siphophage)** | T4P, variable | Electron microscopy | *Pae* | [2] |
| **C5 (podophage)** | PilA | Electron microscopy | *Pae* | [2] |
| D3112 (siphophage) | T4P | T4P mutants | *Pae* | [3] |
| **DLP1 (siphophage)** | PilA | Electron microscopy | *Pae, S. maltophilia* | [4] |
| DMS3 (siphophage) | PilA | Electron microscopy | *Pae* | [5], this study |
| **F116 (siphophage)** | PilA | Electron microscopy | *Pae* | [6] |
| JBD68 (siphophage) | T4P Tip | Electron microscopy | *Pae* | This study |
| Les2 (siphophage) | T4P | T4P mutants | *Pae* | This study |
| Les3 (siphophage) | T4P | T4P mutants | *Pae* | This study |
| LUZ19 (podophage) | T4P | T4P mutants | *Pae* | [7] |
| **M6 (siphophage)** | PilA | Electron microscopy | *Pae* | [2] |
| **PE69 (podophage)** | PilA | Electron microscopy | *Pae* | [2] |
| **PO4 (siphophage)** | PilA | Electron microscopy | *Pae* | [8,9] |
| **PP7 (RNA)** | PilA | Electron microscopy | *Pae* | [10] |
| **PRR1 (RNA)** | PilA | Electron microscopy | *Pae* | [11] |
| **vB_Pae_QDWS (podophage)** | PilA | Electron microscopy | *Pae* | [12] |
| **vB_Pae_Tr (siphophage)** | PilA | Electron microscopy | *Pae* | [13] |
| **vB_Pae_W3 (siphophage)** | PilA | Electron microscopy | *Pae* | [12] |
| ΦKMV (podophage) | T4P | T4P mutants | *Pae* | [14] |
| ΦKZ (jumbo) | T4P | T4P mutants | *Pae* | [15,16] |

**Supplementary Table 1| Phages previously shown to interact with the T4P.** Phages previously shown to interact with PilA using electron microscopy (in bold) and the T4P-dependent phages used in this study.

| Protein Accession | Protein Length (aa) | Phage Name | Host Genus | Phage Genome Size (kbp) | Phage type |
|---|---|---|---|---|---|
| YP_004306725.1 | 59 | KPP10 | *Pseudomonas* | 88.3 | Nankokuvirus |
| CEF89520.1 | 65 | Ab17 | *Pseudomonas* | 83.5 | Nankokuvirus |
| CEQ38340.1 | 59 | Av06 | *Pseudomonas* | 84.7 | Nankokuvirus |
| QIQ63960.1 | 59 | Epa24 | *Pseudomonas* | 88.7 | Nankokuvirus |
| YP_009124427.1 | 62 | Ab03 | *Pseudomonas* | 86.2 | Nankokuvirus |
| YP_009604653.1 | 59 | G1 | *Pseudomonas* | 87.6 | Nankokuvirus |
| YP_009126540.1 | 157 | VpKK5 | *Vibrio* | 56.6 | Queuovirinae |
| UZT28669.1 | 155 | 033B | *Vibrio* | 55.8 | Queuovirinae |
| UOX40479.1 | 152 | AhFM11 | *Aeromonas* | 168 | Biquartavirus |
| QBX32783.1 | 150 | Asfd_1 | *Aeromonas* | 169 | Biquartavirus |
| UYD57450.1 | 151 | B614 | *Aeromonas* | 170 | Biquartavirus |
| YP_009134460.1 | 127 | ACG-2014f | *Synechococcus* | 228 | Atlauavirus |
| QLF86085.1 | 143 | S-CAM7 | *Synechococcus* | 221 | Atlauavirus |

**Supplementary Table 2| PilA-related proteins encoded in various phages.**

| Phage | Gene | Accession Number |
|---|---|---|
| JBD68 | P-FimU | ARM70472 |
| F10 | P-FimU | YP 001293355 |
| Les2 | P-FimU | WP 012613587.1 |
| Les3 | P-FimU | WP 012613709.1 |
| BWH | P-FimU | ASC96364.1 |
| CAAD | P-FimU | WP 049950365.1 |
| NSYG | P-FimU | WP 124176026.1 |
| JBD68 | RBP | ARM70481.1 |
| F10 | RBP | YP 001293364.1 |
| Les2 | RBP | WP 010791961 |
| Les3 | RBP | WP 012613716.1 |
| BWH | RBP | ASC96387.1 |
| CAAD | RBP | WP 226342069.1 |
| NSYG | RBP | WP 124176025.1 |
| JBD68 | Gp24 | ARM70486.1 |
| F10 | Gp24 | YP 001293369.1 |
| Les2 | Gp24 | WP 071536897.1 |
| Les3 | Gp24 | WP 012613719.1 |
| BWH | Gp24 | ASC96392.1 |

| Strain | Gene | Accession Number |
|---|---|---|
| *P. aeruginosa* PAO1 | FimU | NP 253240.1 |
| *X. fastidiosa* M23 | PilA | WP 012382763.1 |
| *X. fastidiosa* GB514 | P-PilA (myo) | YP 006001458.1 |
| *X. fastidiosa* M23 | P-PilA (myo) | YP 001829845.1 |
| *X. fastidiosa* Temecula | P-PilA (myo) | NP 779285.1 |
| *X. fastidiosa* 9a5c | P-PilA (sipho) | NP 297777.1 |
| *X. fastidiosa* M12 | P-PilA (podo) | YP 001775510.1 |

**Supplementary Table 3| Relevant protein accession numbers.**

| Bacterial Strains | Details | Source |
|---|---|---|
| *Pseudomonas aeruginosa* | | |
| PAO1 | Wild-type | Lab collection |
| PAK | Wildt-type | Lab collection |
| PAO1 ΔfimU | fimU mutant, ISlacZ/hah transposon insertion | [17] |
| PAO1 ΔpilA | pilA mutant, ISlacZ/hah transposon insertion | [17] |
| PAO1 ΔpilT | pilT mutant, ISlacZ/hah transposon insertion | [17] |
| PAO1 I-C CRISPR | PAO1 with integrated I-C CRISPR system | [18] |
| PAO1 I-C CRISPR Δ*hel* | PAO1 with integrated I-C CRISPR system with mutated helicase | [18] |
| *Escherichia coli* | | |
| DH5α | | Lab collection |
| SM10 λpir | For conjugation of Pex18Gm into *P. aeruginosa* | Lab collection |
| **Bacteriophages** | | |
| JBD68 | | Lab collection, [19] |
| JBD68 Δp-fimU | JBD68 with 114 bp deletion in the gene eNcoding P-FimU generated using I-C CRISPR-Cas system | This study |
| JBD68 Δrbp | JBD68 with an in-frame, internal deletion in the gene eNcoding RBP (amino acids Δ80-380) | This study |
| JBD68 Δ24 | JBD68 with 285 bp deletion in the gene eNcoding Gp24 generated using I-C CRISPR-Cas system | Lab collection |
| F10 | | Lab collection, [20,21] |
| Les2 | Isolated from *P. aeruginosa* LESB58 | Lab collection, [22] |
| Les3 | Isolated from *P. aeruginosa* LESB58 | Lab collection, [22] |
| Les3 Δ*p-fimU* | Les3 with 356 bp deletion in the gene eNcoding P-FimU generated using I-C CRISPR | This study |
| DMS3 | | Lab collection |
| D3112 | | Lab collection |
| **Plasmids** | | |
| pHERD30T | Arabinose inducible broad host range vector, aacC1 marker | [23] |
| Pex18Gm | *E. coli - P. aeruginosa* conjugation vector, aacC1 marker, SacB | [24] |
| PAO1 FimU | PAO1 fimU cloned into the MCS of pHERD30T using NcoI and HindIII | This study |
| PA14 FimU | PA14 fimU cloned into the MCS of pHERD30T using NcoI and HindIII | This study |
| JBD68 P-FimU | JBD68 p-fimU cloned into the MCS of pHERD30T using NcoI and HindIII | This study |
| Les P-FimU | Les3 p-fimU cloned into the MCS of pHERD30T using NcoI and HindIII | This study |
| F10 P-FimU | F10 p-fimU cloned into the MCS of pHERD30T using NcoI and HindIII | This study |
| BWH P-FimU | BWH p-fimU cloned into the MCS of pHERD30T using NcoI and HindIII | This study |
| CAAD P-FimU | CAAD p-fimU cloned into the MCS of pHERD30T using NcoI and HindIII | This study |
| NSYG P-FimU | NSYG p-fimU cloned into the MCS of pHERD30T using NcoI and HindIII | This study |
| Gp61 | *61* of JBD26 cloned into the MCS of pHERD30T with an N-terminal 6xHis-tag | K.Maxwell lab |
| $RBP_{JBD68}$ | ARM70480.1 & ARM70481.1 cloned into the MCS of pHERD30T using EcoRI and HindIII | This study |
| $RBP_{Les2}$ | WP_010791962.1 & WP_010791961 cloned into the MCS of pHERD30T using EcoRI and KpnI | This study |

**Supplementary Table 4| Strains, bacteriophages and plasmids used in this study.**

| Primer | Sequence | Restriction enzyme |
|---|---|---|
| PAO1 *fimU* F | **AAACCATGGCC**TCATATCGTTCCAACTCGACCGG | NcoI |
| PAO1 *fimU* R | **AAAAAGCTTT**CAATAGCATGACTGGGGCGC | HindIII |
| PA14 *fimU* F | **AAACCATGGCC**CGCTCTATTTGTCGCAGCGCC | NcoI |
| PA14 *fimU* R | **AAAAAGCTTT**CAGTTACAGCTGTCCGGTTGCTTTG | HindIII |
| JBD68 *p-fimU* F | **AAACCATGGCC**TACTCTAGGTCGCGCGGATTTACC | NcoI |
| JBD68 *p-fimU* R | **AAAAAGCTTT**CAGCAGCCAGAAGCGTCCC | HindIII |
| F10 *p-fimU* F | **AAACCATGGCC**TACTCTAGGTCGCGCGGATTTA | NcoI |
| F10 *p-fimU* R | **AAAAAGCTT**CTAACAGTCAAAAGTATCCCAGGATTCC | HindIII |
| Les3 *p-fimU* F | **AAACCATGGCC**TACTCTAGGTCGCGCGGATTTACC | NcoI |
| Les3 *p-fimU* R | **AAAAAGCTTT**CAGCAGCCAGAAGTGTCCCA | HindIII |
| BWH *p-fimU* F | **AAACCATGG**CCGGCTCCAAG | NcoI |
| BWH *p-fimU* R | **AAAAAGCTT**CTAACTGGGTTTTACAGCGCAGGc | HindIII |
| CAAD *p-fimU* F | **AAACCATGG**GTTGAAATGTACTCTAGGTCGCGCG | NcoI |
| CAAD *p-fimU* R | **AAAAAGCTTT**CAGGTTGGTTTCTTGGCGCA | HindIII |
| NSYG *p-fimU* F | **AAACCATGG**ATTCGCATAAATCAAGCTCAGCGTGG | NcoI |
| NSYG *p-fimU* R | **AAAAAGCTT**CTATTGGTCCTTGGCACACTGTG | HindIII |
| JBD68 *rbp* F | **AAAGAATTC**ATGGCTCTTTCAGATGAGCGC | EcoRI |
| JBD68 *rbp* R | **AAAAAGCTTT**CAAACGTAGGAATAGAAAGCGTTCG | HindIII |
| Les2 *rbp* F | **AAAGAATTC**ATGGCTCTATCAGATGAGCGC | EcoRI |
| Les2 *rbp* R | **AAAGGTACC**TCAAACGTAGGAATAGAAAGCGCT | KpnI |
| Les3 *p-fimU* I-C spacer F | **GAAAC**TGGCTGTGTTGTGTTTGCTTACAATGGTTCAATA**G** | BsaI |
| Les3 *p-fimU* I-C spacer R | **GCGAC**TATTGAACCATTGTAAGCAAACACAACACAGCCA**G** | BsaI |
| JBD68 *p-fimU* I-C spacer F | **GAAAC**CGTCGTTTGTGAATCTCATAAAAGGCAATAGCAT**G** | BsaI |
| JBD68 *p-fimU* I-C spacer R | **GCGAC**ATGCTATTGCCTTTTATGAGATTCACAAACGACG**G** | BsaI |
| JBD68 *24* I-C spacer F | **GAAAC**CGTCGTTTGTGAATCTCATAAAAGGCAATAGCAT**G** | BsaI |
| JBD68 *24* I-C spacer R | **GCGAC**ATGCTATTGCCTTTTATGAGATTCACAAACGACG**G** | BsaI |

**Supplementary Table 5| Oligonucleotides used in this study.**

### Supplemental Table References
